# Sleep and circadian rhythm disruption alters the lung transcriptome to predispose to viral infection

**DOI:** 10.1101/2022.02.28.482377

**Authors:** Lewis Taylor, Felix Von Lendenfeld, Anna Ashton, Harshmeena Sanghani, Eric Tam, Laura Usselmann, Maria Veretennikova, Robert Dallmann, Jane A McKeating, Sridhar Vasudevan, Aarti Jagannath

## Abstract

Sleep and circadian rhythm disruption (SCRD), as encountered during shift work, increases the risk of respiratory viral infection including SARS-CoV-2. However, the mechanism(s) underpinning higher rates of respiratory viral infection following SCRD remain poorly characterised. To address this, we investigated the effects of acute sleep deprivation on the mouse lung transcriptome. Here we show that sleep deprivation profoundly alters the transcriptional landscape of the lung, causing the suppression of both innate and adaptive immune systems, disrupting the circadian clock, and activating genes implicated in SARS-CoV-2 replication, thereby generating a lung environment that promotes viral infection and associated disease pathogenesis. Our study provides a mechanistic explanation of how SCRD increases the risk of respiratory viral infections including SARS-CoV-2 and highlights therapeutic avenues for the prevention and treatment of COVID-19.

## INTRODUCTION

Respiratory viral infections are among the leading causes of mortality worldwide and present a global medical and economic challenge ^1,2^. Each year, billions of infections lead to millions of deaths, with the annual financial burden estimated at over $100 billion in the United States alone ^3,4^. The recent emergence of severe acute respiratory syndrome coronavirus type 2 (SARS-CoV-2), the causative agent of COVID-19 ^5^, has highlighted the impact of respiratory viral infections, with more than 400 million SARS-CoV-2 infections and 5.7 million COVID-19 deaths to date (https://coronavirus.jhu.edu/map.html). An increased understanding of the risk factors and mechanisms driving severe respiratory disease will inform new treatment options. Sleep and circadian disruption have been reported to cause an increased risk of respiratory infections in mice and humans ^6–10^ and accumulating evidence suggests that shift work and the associated sleep deprivation and circadian rhythm misalignment are risk factors for COVID-19 ^11–16^. Yet, a mechanistic explanation of how sleep and circadian disruption causes higher rates of viral infections remains to be determined.

The immune system is under tight sleep and circadian control. The circadian clock, a molecular transcriptional/translational feedback loop capable of aligning to the external day/night cycle ^17,18^, generates circadian rhythms; 24-hour oscillations in physiology and behaviour such as hormone secretion, metabolism, sleep, and immune function ^19^. Indeed, leukocyte trafficking, host-pathogen interaction, and immune cell activation all display diurnal rhythms ^20^. Furthermore, circadian differences in immune responses to vaccination, as well as a diverse range of pathogens and pathogen-derived products are well documented ^21,22^. Immune responses to Influenza A, Hepatitis A and SARS-CoV-2 vaccines ^23–25^, and the infectivity of multiple viruses, including Influenza, is dependent on the time of virus challenge ^26–29^. Disrupting the circadian system in experimental model systems has been reported to increase pro-inflammatory cytokine levels ^30^, perturb immune cell function and trafficking ^31^. Furthermore, it can promote the replication of a wide range of clinically important viruses including hepatitis B and C, Parainfluenza Virus Type 3, Respiratory Syncytial and Influenza A viruses ^6,27,32–34^, emphasising a central role of the circadian clock in regulating viral infection ^35^.

Sleep is one of the most essential circadian regulated behaviours; however, sleep and its homeostasis can be modified and disrupted independently from the circadian clock ^36,37^. Sleep disruption also leads to immune dysfunction, reducing natural killer cell activity ^38^, modifying pro-inflammatory cytokine production ^39–42^ and blood leukocyte numbers ^43^. Importantly, sleep disruption impairs circadian and immune gene expression in multiple tissues ^44^, including the mouse brain ^45^, liver ^46^, and lung ^47,48^. A similar disruption of the circadian clock and immune system is seen in blood samples from sleep deprived human subjects ^49–51^. This dual sleep and circadian rhythm disruption (SCRD) is often encountered by shift workers, particularly those working at night, and is a well-established risk factor for respiratory viral infections. The common cold ^52^, Influenza ^6,7,26^, and indeed upper respiratory viral infections in general ^8,10^ are all significantly increased following SCRD. The recent global research effort on SARS-CoV-2 has resulted in multiple studies reporting an association between shift work, sleep disruption and the risk of developing severe COVID-19 ^12,14–16,53^.

Despite the increased risk of respiratory viral infections in shift workers, and the established links between sleep, circadian rhythmicity and immune function, the molecular mechanism(s) underpinning higher rates of viral infection following SCRD remain poorly characterised. We investigated the effects of acute sleep deprivation on the mouse lung transcriptome and perturbation of host pathways recently identified to regulate SARS-CoV-2 infection ^54–57^. Here we show that 6 hours of sleep deprivation in mice profoundly alters the transcriptional architecture of the lung, with a majority of differentially expressed genes associated with host pathways that are essential for viral replication and a suppression of immune and circadian regulated genes with blunted circadian rhythmicity. Moreover, we found that SD causes the differential expression of several host factors implicated in SARS-CoV-2 infection, likely impacting SARS-CoV-2 entry, replication, and trafficking. Together, these data suggest that sleep deprivation alters the lung to provide an environment that promotes respiratory viral infection and pathogenesis.

## RESULTS AND DISCUSSION

### Acute sleep deprivation alters the lung transcriptome and dampens immune-associated gene expression

To assess the effect of acute sleep deprivation on the lung transcriptome, RNA sequencing (RNA-Seq) was performed on lung tissue isolated from control or six-hour sleep deprived (SD) C57BL/6 mice (Fig. 1a). Gene expression analysis identified 2,366 upregulated and and 2,157 downregulated following SD (Fig. 1b, c). We validated our RNA-Seq dataset using qRT-PCR and independent SD lung samples and observed correlated results for several top differential genes and clock genes, confirming the robustness of our transcriptomic analysis (Supplementary Fig. 1). Gene ontology (GO) biological pathway (BP) enrichment analysis of SD upregulated genes showed an enrichment in signal transduction (kinase activity and response to steroid hormones), as well as generic biological processes that are also implicated in viral entry and RNA replication, such as autophagy, Golgi organization, and cellular protein localization (Fig. 1d). Similar results were observed with Kyoto Encyclopaedia of Genes and Genomes (KEGG) pathway analysis of SD upregulated genes, highlighting protein processing in ER, autophagy, and endocytosis (Fig. 1e). We noted an enrichment for circadian rhythm genes (Fig. 1e). Analysis of the SD downregulated genes showed a repression of immune system pathways, including lymphocyte differentiation and proliferation, leukocyte migration, and cytokine production, such as TNF-α, IL-1 and IFN-γ production (Fig. 1f). KEGG pathway analysis displayed a similar enrichment for immune associated terms in the SD downregulated gene population (Fig. 1g). Interestingly, *Furin*, which cleaves the SARS-CoV-2 spike protein and regulates particle entry ^58^, was upregulated following SD, whilst several Toll-like receptors (TLRs), which initiate innate immune responses, were all downregulated, including *Tlr3, Tlr7*, and *Tlr9* that have been shown to regulate COVID-19 pathogenesis ^59,60^ (Fig. 1h). Alongside, the levels of *Nfkbia*, a major negative regulator of pro-inflammatory transcription factor nuclear factor kappa (NF-κB), and its repressor, *Tle1*, were increased by 46% and 43% after SD, respectively. Furthermore, GO BP analysis revealed another twelve SD upregulated genes implicated in negative regulation of NF-κB, and 23 SD downregulated genes encoding positive regulators of NF-κB, with the net effect being increased NF-κB function. In total, 215 significant GO BP terms were enriched for amongst the SD downregulated genes (adjusted p value < 0.01), of which 154 (72%) comprised immune system pathways. Of these, innate and adaptive immunity specific terms comprised 18% and 23% respectively (Fig. 1i). Together, these findings suggest that in the lung, acute SD suppresses both the innate and adaptive immune system and impacts multiple pathways important for viral host cell entry, intracellular replication, and trafficking.

**Fig. 1.**
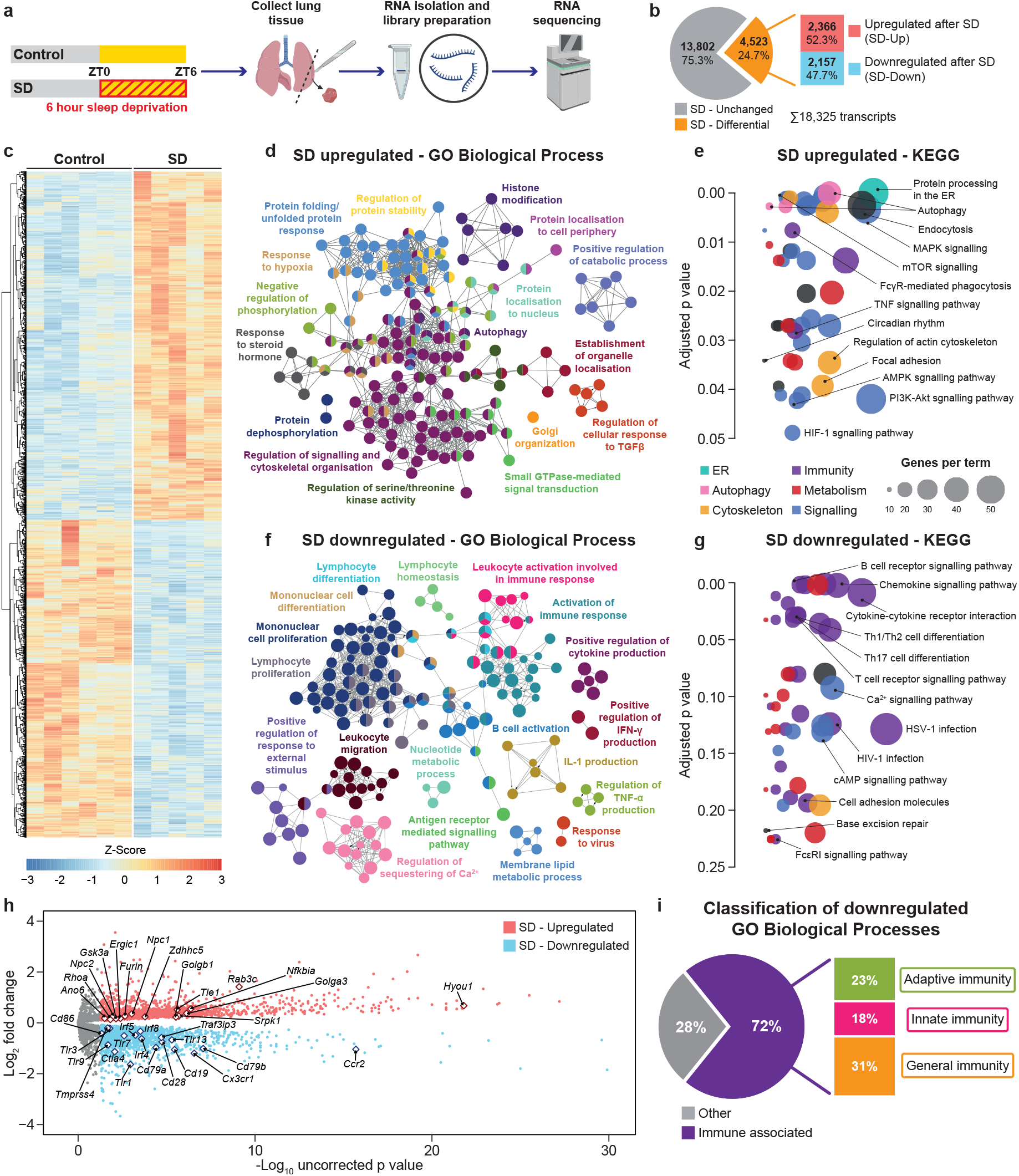
Acute sleep deprivation profoundly alters the lung transcriptome, dampening the immune system and upregulating pathways involved in viral infectivity. **a** WT animals were allowed to sleep *ad libitum* (Control), or sleep deprived (SD) between ZT0 – ZT6. Lung tissue was collected and subjected to RNA-sequencing. **b** Of the 18,325 transcripts identified, 4,523 were differentially expressed following SD, with 2,366 upregulated (SD-Up) and 2,157 downregulated (SD-Down). **c** Heatmap of SD differential genes. **d** GO Biological Process (BP) enrichment analysis and term network visualisation, and **e** KEGG pathway enrichment analysis of SD upregulated genes. **f** GO BP enrichment analysis and term network visualisation, and **g** KEGG pathway enrichment analysis of SD downregulated genes. GO BP/functional grouping is indicated by colour, number of terms/genes is indicated by node size, and edges reflect the relationships between the GO BP terms. **h** Volcano plot of the SD differential genes, with SD-Up (red) and SD-Down (blue) genes highlighted. **i** Classification of the SD downregulated GO BP terms found that 72% were immune associated, and of these 23% were terms involving adaptive immunity, 18% innate immunity and 31% general immunity. For **c** data are Z-score normalised per row. n=5-6. Statistical analysis of RNA sequencing data was conducted using DESeq2, and genes with an adjusted p value of < 0.05 were considered significant.

### Acute sleep deprivation dysregulates the circadian system in the lung

Acute SD was previously reported to disrupt circadian rhythmic gene expression in multiple peripheral tissues ^44,47,48,61^, therefore we examined the full extent of sleep deprivation on circadian processes in the lung. Rhythmic genes in the mouse lung were identified by sequencing the lung transcriptome at four time points throughout the day separated by 6-hour intervals (Zeitgeber time (ZT)2, ZT8, ZT14, and ZT20). We identified 2,029 significantly cycling genes in the mouse lung with a 24-hour period (JTK q-value < 0.05) (Fig. 2a). Interestingly, of these significantly cycling genes, 911 were also disrupted by SD, highlighting that almost 50% of rhythmic genes in the lung are SD sensitive (Fig. 2b). GO BP enrichment analysis of the 3,532 genes that were non-rhythmic, but SD-differential, revealed immune system associated terms such as leukocyte activation and migration (Fig. 2c), indicating that many of these immune genes were not circadian regulated, and instead were directly impacted by SD. Notably however, GO BP analysis of genes that were both rhythmic and SD-differential showed an enrichment for circadian regulation of gene expression, demonstrating that SD alters the circadian regulatory landscape of the lung (Fig. 2d). Indeed, pathways regulating metabolism, signalling, RNA processing, protein folding, and post-translational protein modification were also rhythmic and dysregulated following SD (Fig. 2d), suggesting a widespread disruption of normal circadian lung physiology.

**Fig. 2.**
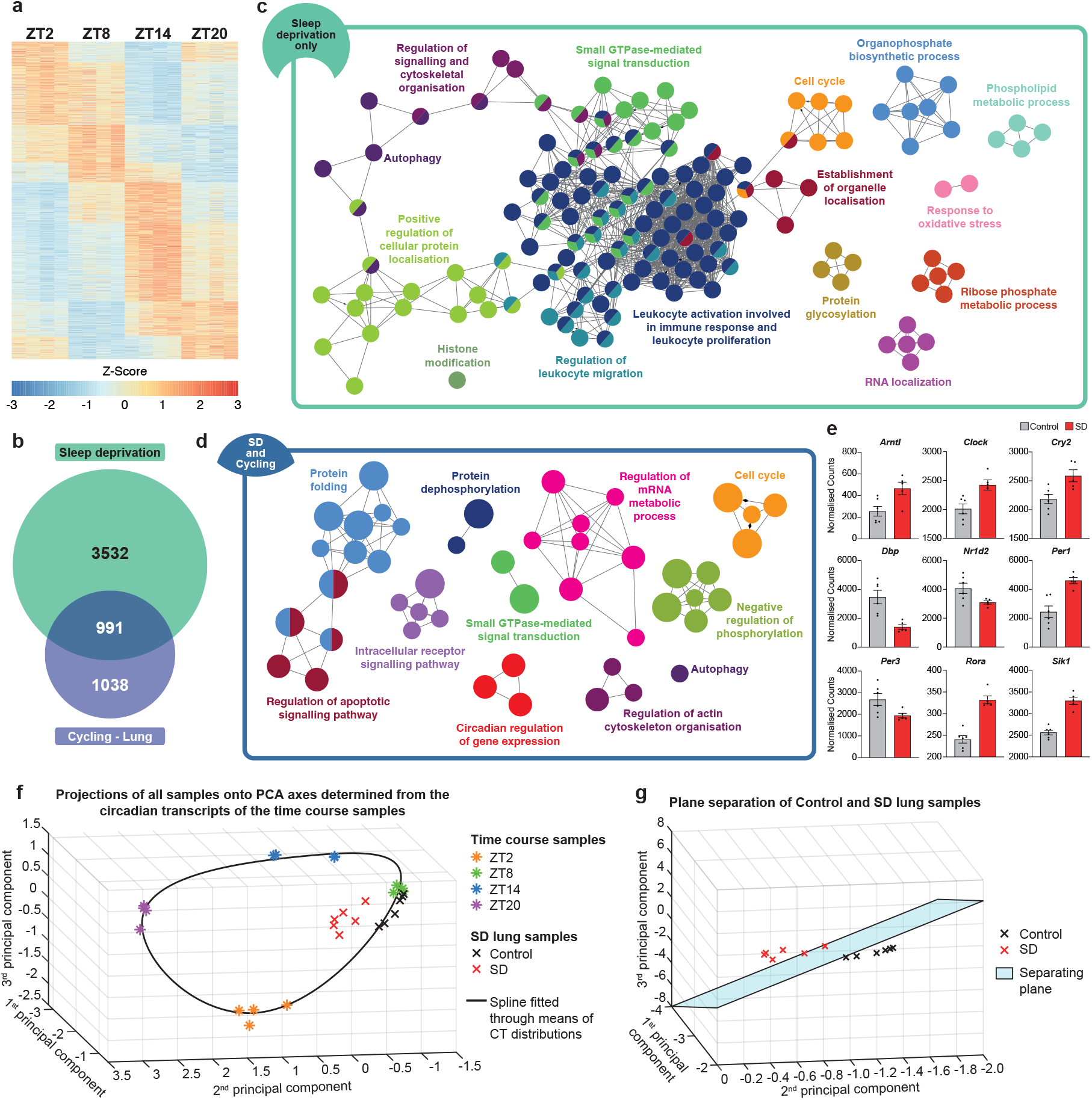
Acute sleep deprivation leads to dysregulation of the circadian system in the lung. WT animals were stably entrained to a 12:12 LD cycle, and then placed into constant darkness. Lung samples were then collected at CT2, CT8, CT14 and CT20 and RNA sequencing conducted. **a** Heatmap of the 2,029 significantly cycling genes in the lung. **b** A comparison with genes differentially expressed following SD with the lung circadian genes found 991 rhythmic genes that were also disrupted by SD. Network visualisation of the significantly enriched GO BP terms of the **c** 3,532 genes that are non-cyclic in the mouse lung, but disrupted by SD (green), and **d** the 991 genes that are cycling in the mouse lung and disrupted by SD (blue). Each node represents a GO BP term. Related terms are grouped by colour and edges reflect the relationship between them. **e** RNA sequencing counts of core circadian clock genes in the control and SD lung samples. **f** PCA projection of the circadian (CT) samples in the PC space determined from 10 known circadian transcripts. The black spline represents the estimated circadian behaviour of mouse lung under constant conditions, and the graph is oriented such that the separation between the CT samples is as clear as possible. The control samples (black crosses) projected near to the black spline at the approximate expected location, however the SD samples (red crosses) did not project to the same location, demonstrating that SD resulted in circadian disruption in the lung. **g** A Support Vector Machine (SVM) approach with the linear kernel was used to find the plane which optimally separated the control and SD samples in the 3D principal component space. The samples were projected onto the normal of this plane, and a clear separation between the two groups can be seen, which was statistically significant (Wilcoxon rank sum test – p = 0.0022). Therefore, SD results in circadian disruption of the lung transcriptome. For **a** data are Z-score normalised per row and for **e** data are mean ± SEM. n=5-6. Cycling genes were determined using MetaCycle, and genes with a corrected q value of < 0.05 were considered as significantly rhythmic. For **f** and **g** statistical analysis was conducted using the Wilcoxon rank sum test.

Therefore, we sought to quantify the degree of circadian rhythm disruption in the lung caused by acute SD. Several circadian transcripts were altered following SD; these included Bmal1 (*Arntl1*), *Clock, Per1, Cry2*, and *Rora* (Fig. 2e), suggesting that at this point of time (ZT6, post SD), the integrity of the core molecular circadian clock, and clock-controlled gene expression, was likely to be compromised. Principal component analysis of the lung transcriptome allowed us to assess the circadian dysfunction in the lung. The principal directions of a group of 10 circadian transcripts from our ZT transcriptomic dataset (Fig. 2a) were used to project all lung samples onto a 3D space and a spline fitted to represent the expected circadian time and behaviour of the lung (Fig. 2f). Any deviation from this spline would represent an abnormal circadian landscape, and indeed, this is what we found. In contrast to the control samples (black crosses), which fell onto the spline in the expected location, the SD samples (red crosses) were displaced, demonstrating that SD disrupted circadian networks in the lung (Fig. 2f). To quantify the impact on the circadian transcriptome, we used a Support Vector Machine approach to locate the plane that maximally separated the control and SD samples. As can be seen in figure 2g, the optimal plane allowed a clear and significant separation between the SD and control groups (Wilcoxon rank sum test p = 0.0022; Fig. 2g). Overall, these data demonstrate that acute SD alters circadian regulation in the lung, and we believe this disruption contributes towards the increased susceptibility to respiratory viral infection.

### Host factors implicated in SARS-CoV-2 infection are differentially expressed in the mouse lung following sleep deprivation

Our data shows that acute SD modifies the transcriptional landscape of the lung in two keys ways to promote infection by respiratory viruses, Firstly, by suppressing the innate and adaptive immune responses, and secondly by disrupting the normal circadian regulatory landscape and physiology of the lung. Three independent studies by Daniloski et al. ^55^, Zhu et al. ^57^, and Wei et al. ^56^ conducted genome-wide CRISPR loss-of-function screens to identify genes regulating SARS-CoV-2 infection. Therefore, we used these data to examine whether SD changes the expression of host factors required by SARS-CoV-2. Daniloski et al. identified 1,200 potentially relevant genes for SARS-CoV-2 replication and investigated the 50 most highly enriched ^55^. Of the 50, 10 were dysregulated following SD (Fig. 3a and d), of which 8 could be assigned a putative function in SARS-CoV-2 replication (Fig. 3g). For example, members of the vacuolar-ATPase proton pump, (ATP6V0B, ATP6AP1, and ATP6V0D1), implicated in activation of the SARS-CoV-2 spike protein that is required for viral entry, and ACTR2 and ACTR3, part of the ARP2/3 complex, which functions in endosomal trafficking pathways. Zhu et al. identified 32 genes with a potential role in viral entry ^57^, of which 8 were SD-differential genes (Fig. 3b and d); four each being up- (NPC1, NPC2, CCDC93, WDR81) and downregulated (COMMD8, COMMD10, ACTR2, ACTR3). All 8 genes play a role in endosomal entry, endolysosomal fusion, or endosome recycling (Fig. 3g). Cross-referencing our SD-differential genes to the 50 most enriched host factors identified by Wei et al. ^56^ revealed an intersection of 11 genes (Fig. 3c), 8 of which were upregulated and associated primarily with transcriptional regulation (DPF2, JMJD6, RAD54L2, CREBBP, RYBP, ELOA, KMT2D, SIK1 – Fig. 3g). The effect of SD on the individual transcripts that encode for these putative SARS-CoV-2 host factors across all three studies is illustrated in figure 3d. Taken together therefore, acute SD clearly amplifies many host factors and processes that influence multiple steps in the SARS-CoV-2 life cycle.

**Fig. 3.**
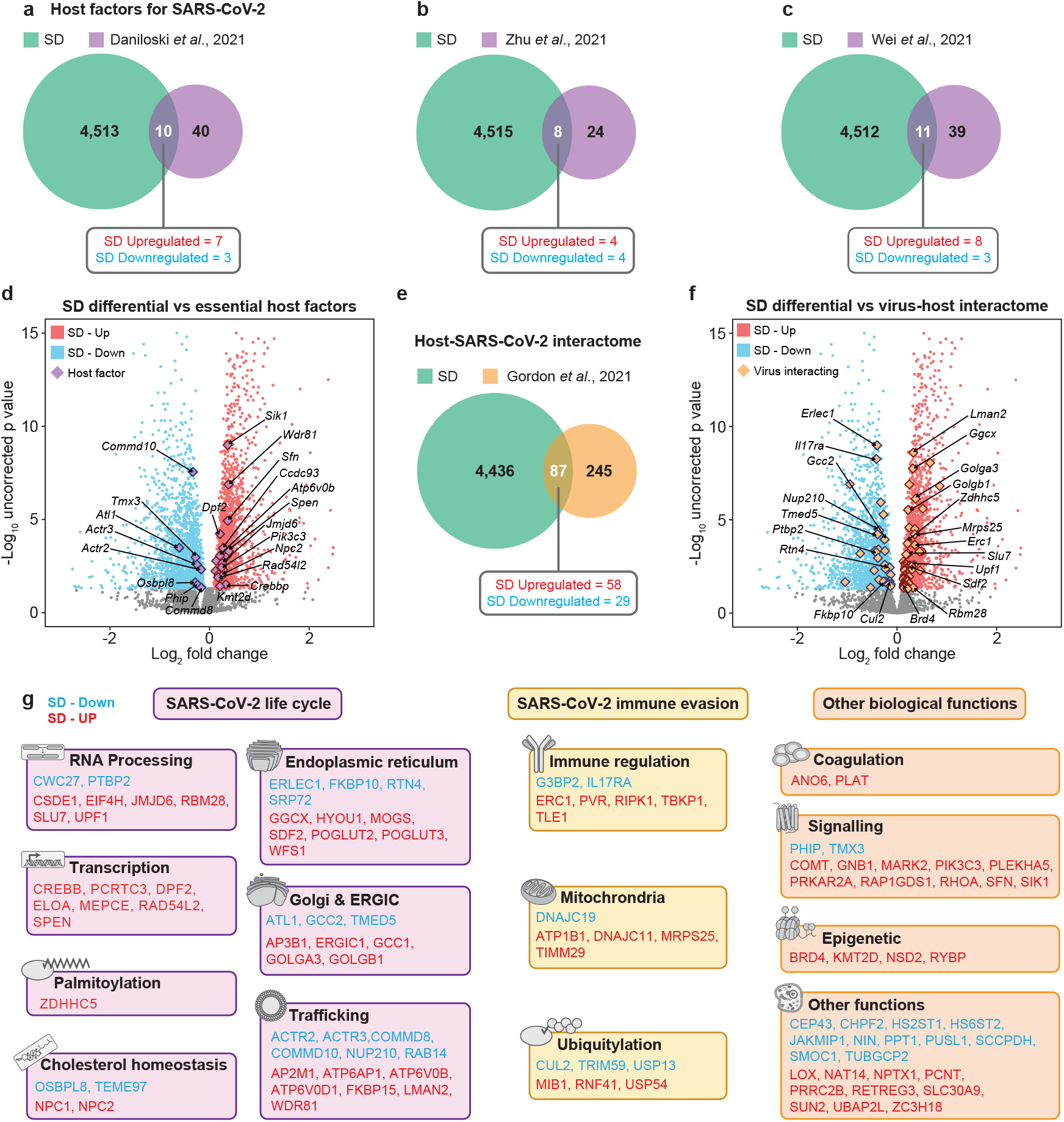
Critical host factors that interact with SARS-CoV-2, and are needed for infection, are differentially expressed in the mouse lung after sleep deprivation. **a**-**c** Venn diagrams of the overlap between all SD differential genes in the mouse lung and critical host factors for viral infection as determined by **a** Daniloski *et al*. (2021), **b** Zhu *et al*. (2021), and **c** Wei *et al*. (2021). **d** Volcano plot of SD differential genes with overlapping critical host factors highlighted (purple diamonds) and a subset labelled. **e** The intersection between SD differential genes in the mouse lung and the SARS-CoV-2-human protein interactome as determined by Gordon *et al*. (2020b). **f** Volcano plot of SD differential genes with overlapping SARS-CoV-2-human protein interactors highlighted (yellow diamonds) and a subset labelled. **g** Functional classification of the critical host factors and the SARS-CoV-2-human protein interactors that were found to be differentially expressed in the mouse lung following SD. Boxes are coloured according to the functional role in viral infectivity. Statistical significance of the overlap between SD differential genes and SARS-CoV-2 host factors and host interactome was assessed by two-tailed Fisher’s exact test. n=5-6. ERGIC = endoplasmic reticulum-Golgi apparatus intermediate compartment.

We next explored if SD-differential genes encode host proteins known to physically interact with SARS-CoV-2 encoded proteins. Gordon et al. interrogated human host factors that interact with 26 of the 29 SARS-CoV-2 proteins ^54^. The authors identified 332 high confidence human-virus protein-protein interactions, of which 87 overlapped with our SD-differential genes (Fig. 3e,f). Interestingly, at least 40 of the overlapping genes have a putative function in viral replication, such as RNA processing, ER protein quality control, or intracellular trafficking (Fig. 3g). Furthermore, 18 of the overlapping host factors are involved in mitochondrial processes, ubiquitination, or immune regulation, that may function in SARS-CoV-2 immune evasion (Fig. 3g). Alongside, regulators of signalling pathways, coagulation, and epigenetic modifiers represent some of the other dysregulated classes of interactors that likely impact SARS-CoV-2 infection. Overall, these findings demonstrate that SD causes the differential expression of several host factors that interact with, and are implicated in, SARS-CoV-2 infection that may potentiate virus replication.

### The effect of sleep deprivation on SARS-CoV-2 life cycle genes

When taken together, our data suggest that acute SD impacts many host processes important for the SARS-CoV-2 life cycle, and we propose a mechanistic pathway, synthesised from the data presented above, by which the SD-differential genes facilitate SARS-CoV-2 entry, replication, and trafficking (Fig. 4). The extracellular transmembrane protease serine 4 (*Tmprss4*), the protease *Furin*, and *Atp6v0b, Atp6ap1*, and *Atp6v0d1* (members of the vacuolar-ATPase proton pump) all contribute towards spike protein activation and cleavage ^62,63^, and were differentially expressed following SD, suggesting an increase in virus entry. Following intracellular capsid uncoating the viral RNA is replicated within double membrane vesicles, translated by host ribosomes, and new virus particles assembled and trafficked via the Golgi/ER pathway for release by exocytosis. All these pathways were dysregulated by SD, including transcriptional modulation, endolysosomal fusion, endosome recycling (Fig. 4).

**Fig. 4.**
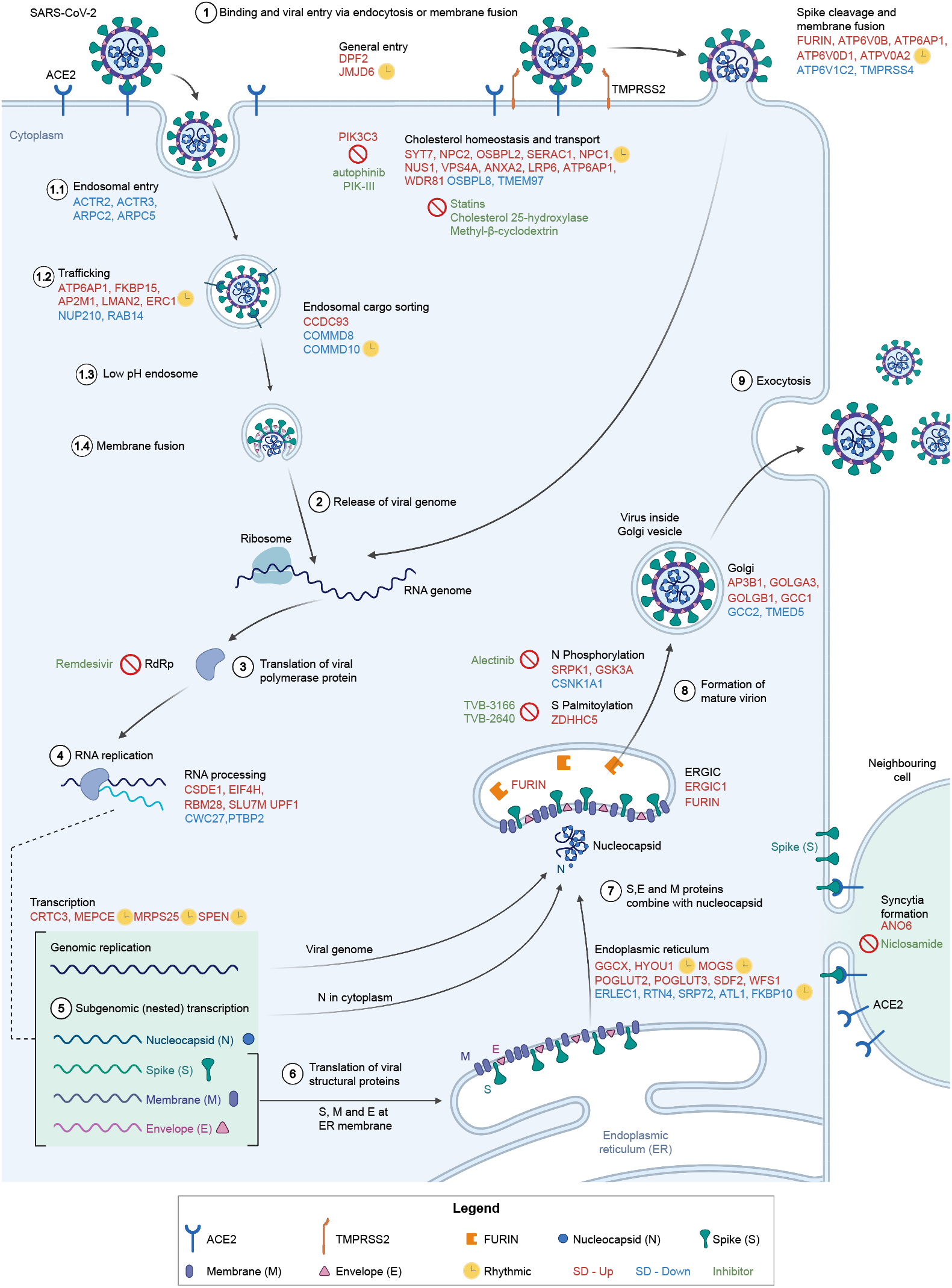
Involvement of differentially expressed genes after sleep deprivation in the SARS-CoV-2 life cycle. (**1**) SARS-CoV-2 binds ACE2 and enters via endocytosis or membrane fusion, depending on the availability of TMPRSS2/4. (**2**) The viral RNA genome is released into the cytoplasm and (**3**-**4**) replicated and (**5**) transcribed by RdRp. (**6**) Viral structural proteins are translated by host ribosomes. (**7**-**8**) The virion assembles and (**9**) is released. All differentially expressed genes shown (red font for SD-Up, blue font for SD-Down) apart from *FURIN, TMPRSS4, GSK3A, SRPK1*, and *CSNK1A1* are critical host factors overlapping with at least one of the studies from Gordon *et al*. (2020b), Daniloski *et al*. (2021), Wei *et al*. (2021), or Zhu *et al*. (2021). Drugs targeting SD differential or viral genes mentioned are in green font. Cycling genes are denoted by a yellow clock. ACE2 = angiotensin-converting enzyme 2, ERGIC = ER-Golgi apparatus intermediate compartment, RdRp = RNA-dependent RNA polymerase, TMPRSS2 = transmembrane protease serine 2. Adapted from Du et al. (2009) and from “Coronavirus Replication Cycle” by BioRender.com. Created with BioRender.com.

Thirteen genes implicated in intracellular cholesterol trafficking (*Tmem97, Syt7, Npc1, Npc2, Osbpl2, Serac1, Nus1, Vps4a, Anxa2, Lrp6, Atp6ap1, Pik3c3*, and *Wdr81*) were differentially expressed following SD, in line with previous findings showing SD driven disruption of cholesterol metabolism ^64^. This was of interest to us, as three of the four cross-referenced SARS-CoV-2 host factor studies (Fig. 3) identified disrupted cholesterol homeostasis as a risk factor for infection. Plasma membrane cholesterol is required for SARS-CoV-2 fusion and cell entry ^65^, a pathway common to most enveloped viruses. Furthermore, statins have been found to reduce recovery time and decrease the risk for COVID-19 morbidity and mortality ^66,67^. How cholesterol impacts SARS-CoV-2 pathogenesis is currently unclear; however, lipid raft disruption, modification of membrane biophysics, alteration of viral stability and maturation, and immune dysfunction have all been suggested as potential mechanisms ^68–70^.

Finally, SD alters post-translational protein modification that regulates multiple aspects of SARS-CoV-2 replication. For example, the viral nucleocapsid protein is phosphorylated by SRPK1, GSK-3a, and CSNK1 ^71^ and genes encoding all three kinases were differentially expressed in the lung after SD. Palmitoylation of the Spike envelope glycoprotein is necessary for infectivity. Knockdown of ZDHHC5, a palmitoyltransferase, resulted in spike protein depalmitoylation and compromised membrane fusion and viral entry ^72^, and SD resulted in increased *Zdhhc5* transcripts in the lung. Overall, these findings suggest that SD could promote SARS-CoV-2 replication by dysregulating many genes involved in its life cycle.

### The effect of sleep deprivation on the anti-SARS-CoV-2 immune response and viral immune evasion

Alongside the impact on viral replication, our data shows that SD can suppress immune associated genes allowing viral persistence. Analysis of the SD lung transcriptome shows altered regulation of several components of the immune system (Fig. 5). The regulators of interferon production, RNF41 and TBKBP1, are targeted by SARS-CoV-2 proteins ^54^ and their genes were differentially expressed following SD. Furthermore, SD caused the differential expression of the E3 ubiquitin ligases *Mib1* and *Trim59*, which induce and repress NF-κB, respectively ^73,74^, alongside the NF-κB repressor, *Tle1*. These proteins have been shown to associate with SARS-CoV-2 proteins, suggesting that infection interferes with the NF-κB pathway as an immune evasion strategy. Accumulating evidence suggests that SARS-CoV-2 exploits the host ubiquitination machinery to evade the innate immune response ^75,76^, and intriguingly, six SD differential genes (*Mib1, Rnf41, Usp54, Cul2*, Trim59, Usp13) functionally implicated in ubiquitination, encode proteins that interact with SARS-CoV-2 proteins ^54^. Severe COVID-19 is sometimes associated with syncytia in the lung; multinucleated single cells formed by the fusion of SARS-CoV-2 infected cells to allow viral genome transfer without activating the immune system ^77^. Recently, ANO6 has been found to regulate syncytia formation ^78^, and interestingly, we found that *Ano6* was upregulated following SD. Finally, mitochondrial manipulation is another approach by which coronaviruses evade the host immune system ^79^, and notably we found five SD differential mitochondrial host genes (*Dnajc19, Atp1b1, Dnajc11, Mrps25*, and *Timm29*) which are known to engage in SARS-CoV-2 protein-protein interactions ^54^. Taken together therefore, this highlights how acute SD may specifically promote immune evasion by SARS-CoV-2.

**Fig. 5.**
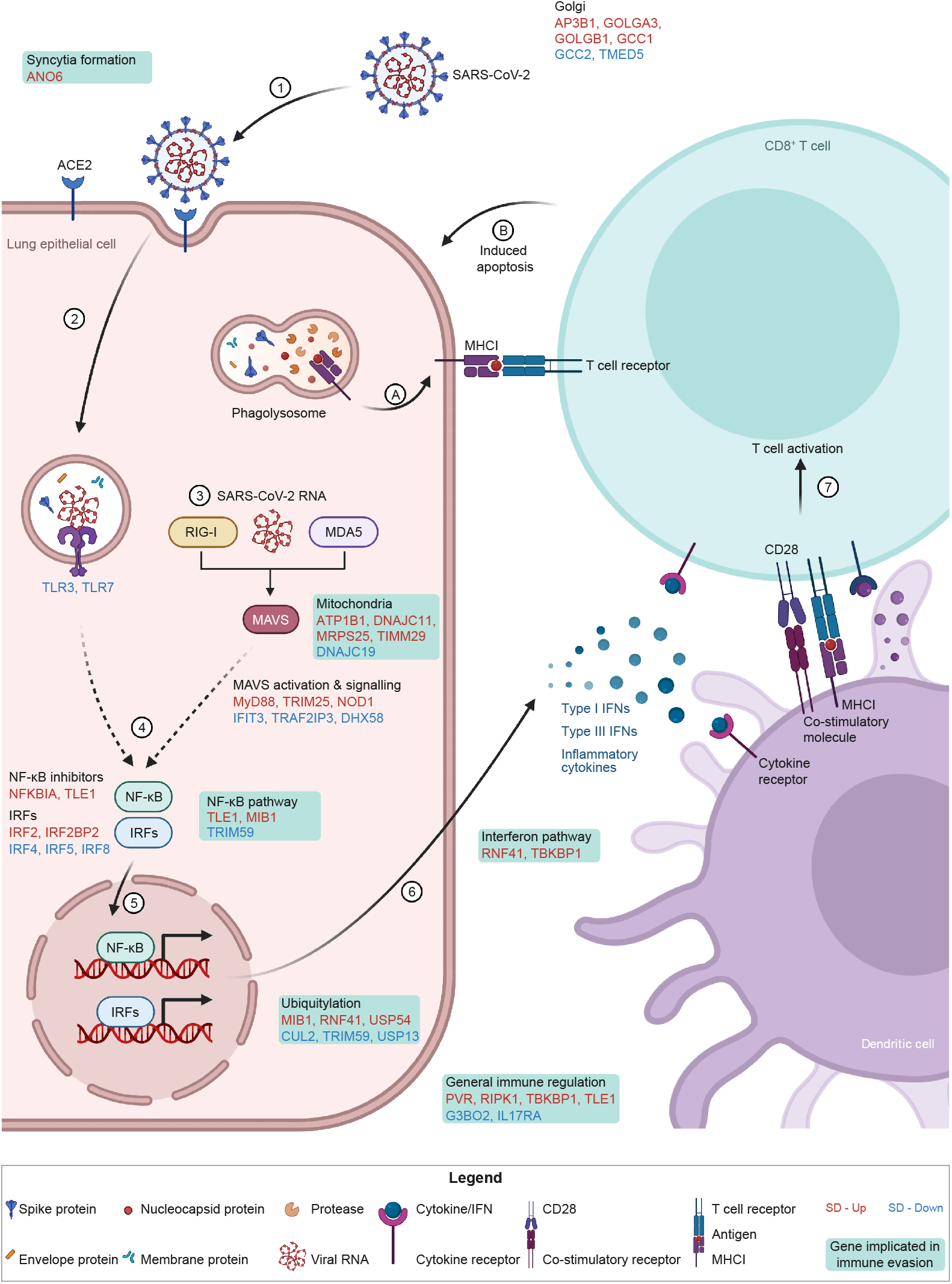
The effect of sleep deprivation on the anti-SARS-CoV-2 immune response and viral immune evasion. (**1**) The virus enters the host cell. Viral RNA is detected by (**2**) endosomal TLRs or (**3**) cytosolic RIG-I and MDA5, which activate MAVS. (**4**)
Both recognition events activate NF-κB and IRFs, which (**5**) translocate into the nucleus to (**6**) drive the expression of IFNs and inflammatory cytokines to amplify the antiviral immune program, for example by priming dendritic cells to sample and display viral antigens to (**7**) activate naive CD8^+^ T cells. Inside the infected host cell, (**A**) viral material is broken down and displayed on the cell surface by MHC class I molecules. If the antigen is recognised by CD8^+^ T cells (**B**) it induces apoptosis of the infected host cell. All differentially expressed genes shown (red font for SD-Up, blue font for SD-Down) are involved in the acute innate immune response against SARS-CoV-2. Genes in green-shaded text boxes are implicated in viral immune evasion, as described in Gordon *et al*., (2020b). IFN = interferon, IRF = interferon regulatory factor, MAVS = mitochondrial antiviral signalling protein, MDA5 = melanoma differentiation-associated protein 5, MHC I = major histocompatibility complex molecule class I, NF-κB = nuclear factor kappa B, RIG-I = retinoic acid-inducible gene 1. Adapted from “Acute Immune Responses to Coronaviruses”, by BioRender.com. Created with BioRender.com.

In conclusion, this study shows that SD alters the transcriptomic landscape in the mouse lung in a manner that could explain the increased risk of respiratory viral infections, as well as severe COVID-19, associated with SCRD and shift work. Suppression of the immune response and promotion of SARS-CoV-2 replication and immune evasion are among the most relevant pathways deregulated by SD. Furthermore, we found a widespread disruption of circadian rhythmicity in the lung following sleep deprivation, which could precipitate and/or exacerbate the negative consequences of SCRD. The hypotheses proposed in this study require validation by challenging mice with SARS-CoV-2 after SD; indeed, this would be an important follow-up study. However, these findings already help explain why SCRD is associated with severe COVID-19 and could guide future efforts towards understanding the mechanisms underlying SARS-CoV-2 pathogenesis. Importantly, our observations are applicable to a wide range of respiratory viruses and may inform avenues to develop new therapeutic efforts.

## ACKNOWLEDGEMENTS

This work was supported by the following sources of funding: BB/N01992X/1 David Phillips fellowship from the BBSRC to AJ. LT is supported by the Oxford-Elysium Health fellowship. JAM is funded by a Wellcome Investigator Award 200838/Z/16/Z, UK Medical Research Council (MRC) project grant MR/R022011/1 and Chinese Academy of Medical Sciences (CAMS) Innovation Fund for Medical Science (CIFMS), China (grant number: 2018-I2M-2-002). LU was funded by UK Medical Research Council Doctoral Training Partnership (MR/N014294/1). MV is supported by a grant from Cancer Research UK (C53720/A29468 to RD).

## AUTHOR CONTRIBUTIONS

LT, FVL, AA, HS, ET and AJ conducted the experiments. LT, FVL, LU, MV and RD analysed data. AJ and SV supervised the study. LT, FVL, JAM and AJ co-wrote and edited the manuscript, with input from all authors.

## DECLARATION OF INTERESTS

The other authors declare no financial interests.

## METHODS

### Animals

All studies were conducted using male C57BL/6 mice over 8 weeks of age and, unless otherwise indicated, animals were group housed with ad libitum access to food and water under a 12:12 hour light/dark cycle (100 lux from white LED lamps). All animal procedures were conducted in accordance with the UK Home Office regulations (Guidance on the Operation of Animals (Scientific Procedures Act) 1986) and the University of Oxford’s Policy on the Use of Animals in Scientific research, following the principles of the 3Rs. For circadian time course analysis, lung tissue was collected at zeitgeber time (ZT)2, ZT8, ZT14, and ZT20. For the sleep deprivation (SD) experiments, animals were kept awake for 6 hours between ZT0 and ZT6 by providing novel objects to elicit exploratory behaviour, as previously described ^80^. The animals were then sacrificed, and lung tissue collected. Control animals were allowed to sleep ad libitum between ZT0 and ZT6.

### RNA extraction and RNA sequencing library preparation

Total RNA from lung tissue samples was extracted using TRIzol and the RNeasy Mini Kit (Qiagen). Lung tissue was mechanically disrupted in 700 μl of TRIzol and 140 μl of chloroform was added and the sample thoroughly mixed. Following a 3 min incubation at RT, the sample was then centrifuged for 15 min at 15,000 xg, 4°C. The clear top layer was then carefully collected, mixed with an equal volume of 70% ethanol and RNA extracted using the RNeasy Mini Kit, with on-column DNase digestion, following the manufacturer’s instructions. RNA was eluted in water and RNA concentration and quality were measured using a TapeStation system (Agilent) with the High Sensitivity RNA ScreenTape assay. mRNA purification and cDNA synthesis for the sequencing library were performed according to the Illumina Stranded mRNA Prep protocol (20040534) using the following index kit: IDT for Illumina RNA UD Indexes Set A, Ligation (20040553). Quality and concentration of the final libraries were checked with the KAPA Library Quantification Kit (Roche Diagnostics) in a StepOnePlus thermal cycler (Applied Biosystems) according to manufacturer’s instructions. All cDNA libraries were sequenced using a paired-end strategy (read length 150 bp) on an Illumina NovaSeq platform.

### qRT-PCR

Total RNA was extracted from mouse lung tissue as detailed above and cDNA was synthesized using the qScript cDNA Synthesis Kit (Quantabio). mRNA was quantified using the QuantiFast SYBR Green PCR Kit (Qiagen) in a StepOnePlus thermal cycler. Cycling conditions were 95 °C for 5 min, and 40 cycles of 95 °C for 10 s, 60 °C for 30 s, 72 °C for 12 s. The cycle thresholds for each gene were normalized using ActB, Gapdh, and Rn18s as housekeeping genes following the 2^^-ΔCt^ method.

### Processing of RNA sequencing data

Raw RNA-Seq data processing (quality control, trimming, mapping to the genome, and read counting) was performed using tools embedded in Galaxy (v21.05) ^81^. The fastqsanger files containing the raw sequencing data were uploaded to the public Galaxy server at usegalaxy.org. FastQC (v0.11.8) (https://www.bioinformatics.babraham.ac.uk/projects/fastqc/) was used for quality control of sequencing data. For quality and adapter trimming, Trim Galore! (v0.6.3) (https://www.bioinformatics.babraham.ac.uk/projects/trim_galore/) was employed to remove low-quality bases, short reads, and Illumina adapters. Nextera transposase was specified as the adapter sequence to be trimmed and Trim Galore! was instructed to remove 1 bp from the 5’ end of both read 1 and 2. FastQC was rerun to assess the quality improvement. High quality reads were then mapped to the Mus musculus (mm10) reference genome using HISAT2 (v2.1.0) ^82^, specifying the strand information as reverse. featureCounts (v2.0.1) ^83^ was run to quantify the number of reads mapped to each gene. The featureCounts built-in mm10 gene annotation file was selected and under paired-end reads options, the option to count fragments instead of reads was enabled. The generated counts files were converted to CSV and downloaded for downstream differential gene expression analysis in R. MultiQC (v1.9) ^84^ was used to aggregate FastQC, HISAT2, and featureCounts results.

### Differential gene expression analysis

To identify differentially expressed genes in the SD and times series (ZT) datasets (adjusted p value < 0.05), the DESeq2 package (v1.32.0) ^85^ was used in R (v4.1.0). Heatmaps were drawn using the pheatmap function from the pheatmap package (v1.0.12). Volcano plots were generated using the ggplot2 package (v.3.3.5). To detect periodicity in the time series (ZT) data, the MetaCycle R package (v1.2.0) was used ^86^. The meta2d function was run using the MetaCycle web application (MetaCycleApp) based on the shiny package (v1.6.0). The following parameters were specified: minper = 24, maxper = 24, ARSdefaultPER = 24, cycMethod = JTK, combinePvalue = fisher. A corrected q value of < 0.05 was considered significantly rhythmic. The MetaCycleApp was downloaded from https://github.com/gangwug/MetaCycleApp.

### Functional enrichment analysis

Functional enrichment analysis of SD-associated genes and cycling genes was conducted using the clusterProfiler R package (v4.0.0) ^87^. GO BP and KEGG analysis was performed using the enrichGO function, with org.Mm.eg.db (v3.13.0) as the Mus musculus genome annotation (GO BP parameters - pvalueCutoff = 0.01, qvalueCutoff = 0.05, pAdjustMethod = Benjamini–Hochberg correction and KEGG paramerters - pvalueCutoff = 0.05). Enriched KEGG terms were visualised using a custom R script. The network interaction between overrepresented GO BP pathways was visualized using the ClueGO application (v2.5.8) ^88^ and its plugin CluePedia (v1.5.8) ^89^ within the desktop version of the Cytoscape software (v3.8.2) ^90^. The yFiles Organic Layout from the yFiles Layout Algorithms application (v1.1.1) ^91^ was used to specify the design.

### Principal component analysis projection of circadian and SD transcript expression

To assess the circadian behaviour of the mouse lung we used principal component analysis (PCA). We first reduced the transcriptomic datasets to 10 circadian features, i. e., transcripts known to be highly rhythmic across murine organ systems (*Arntl, Per2, Per3, Tef, Hlf, Dbp, Nr1d1, Nr1d2, Npas2*, and *Dtx4*) ^92^. The resultant transcript x sample matrices were log-transformed and then Z-score normalised column-wise to prepare the data for dimensionality reduction. Singular value decomposition was applied to the 16 samples collected at times ZT2, ZT8, ZT14, and ZT20 to obtain the principal directions (using the svd function in MATLAB v2020b). All lung samples (time course and SD) were then projected onto the 3D principal component space generated from the first three principal directions of the time course samples. The time point means of the projected time course samples were estimated by fitting Gaussian distributions. A shape-preserving cubic spline was fitted through the estimated means of the projected time course samples to approximate the expected circadian behaviour of the mouse lung (using the csape function in MATLAB). The Support Vector Machine approach (package gensvm v.0.1.5 in R v.4.1.1) with the linear kernel was then used to find the equation of the plane which optimally separated the control and SD lung samples in the 3D principal component space, and then all samples were projected onto the normal of the plane. A Wilcoxon’s rank sum test was carried out in MATLAB (ranksum function) for the projections on the normal to determine whether the null hypothesis that the control and SD samples belonged to the same population (same median) could be rejected.

### Statistical analysis

All data are expressed as mean + or ± SEM, and n represents the number of independent animals or replicates per group, as detailed in each figure legend. For comparisons between two groups only, a one-tailed unpaired Student’s t-test was used. Statistical significance of gene set overlaps was assessed by two-tailed Fisher’s exact test, assuming 21,647 total genes in the lung transcriptome as determined by the RNA-Seq data from SD and time series analysis in this study. Correlation between the qRT-PCR and RNA-Seq expression data was examined using two-tailed Pearson correlation analysis. Statistical testing was performed in R, MATLAB, and GraphPad Prism 9 (v9.1.2).

### Data availability

The authors declare that all data supporting the findings of this study are available within the article and its Supplementary Information files or are available from the authors upon request. RNA sequencing data from this study will be deposited on NCBI with a GEO accession ID.

---

**Supplementary Fig. 1.**
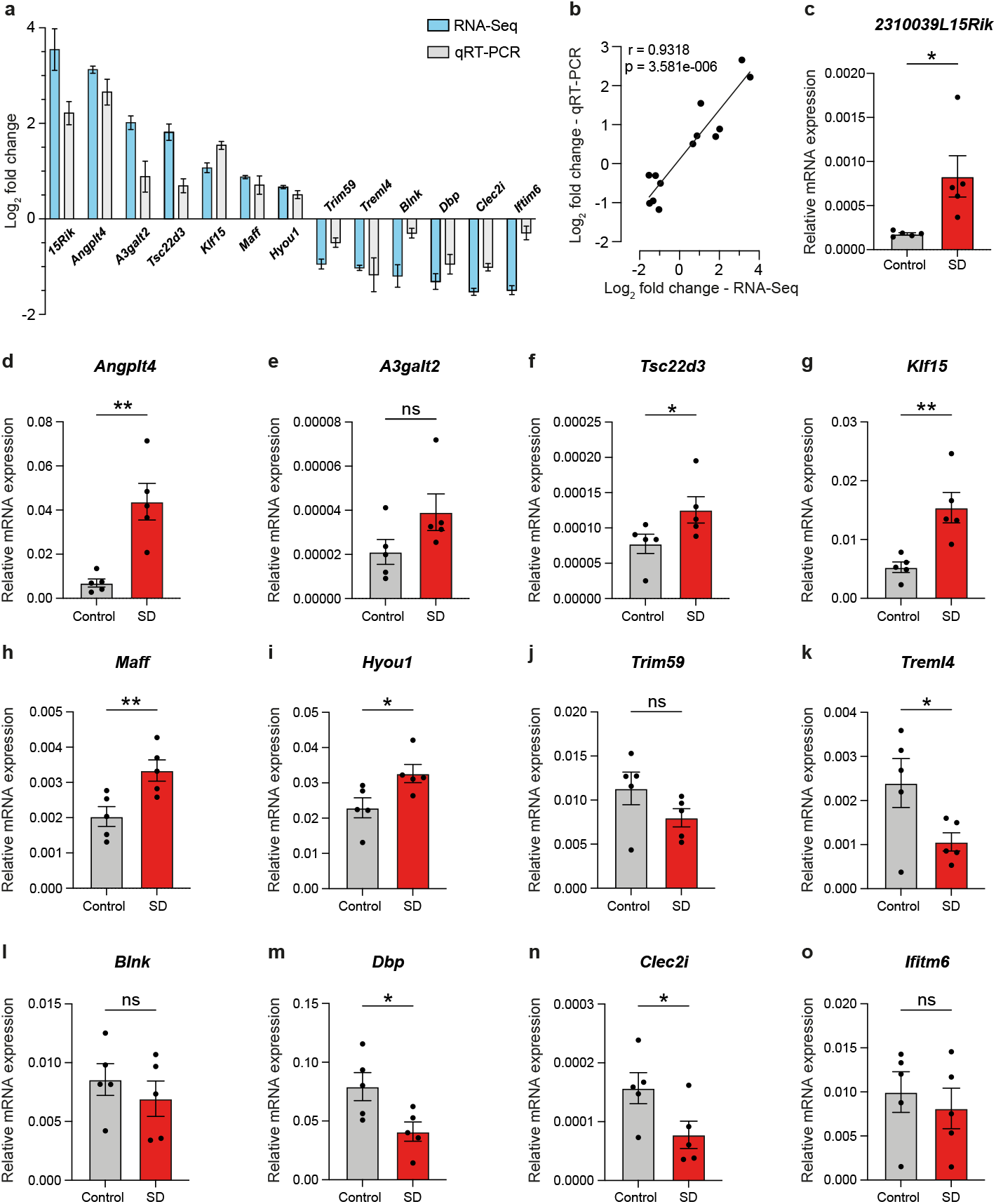
qRT-PCR analysis of independent, sleep deprived lung samples demonstrates the robustness of our RNA sequencing data. Thirteen genes that has differential expression following SD in the lung were selected for qRT-PCR validation using independent control and SD lung samples. **a** Direct comparison of the Log_2_ fold change detected for each gene by RNA-Seq and qRT-PCR. **b** Pearson correlation analysis between RNA-Seq and qRT-PCR gene expression data demonstrates these are significantly positively correlated. Relative expression of **c** *2310039L15Rik*, **d** *Angplt4*, **e** *A3galt2*, **f** *Tsc22d3*, **g** *Klf15*, **h** *Maff*, **i** *Hyou1*, **j** *Trim59*, **k** *Treml4*, **l** *Blnk*, **m** *Dbp*, **n** *Clec2i* and **o** *Ifitm6* in control and SD lung samples. For **b** data are mean, for **a** and **c**-**o** data are mean ± SEM. n=5. Statistical analysis was conducted by unpaired one-tailed student’s t-test. ns P > 0.05, * P < 0.05, ** P < 0.01.

## Notes

### Competing Interest Statement

The authors have declared no competing interest.

